# Measuring drift by mean deviation: unequal breeding sex ratio revisited

**DOI:** 10.1101/2022.02.15.480610

**Authors:** Ben Qin

## Abstract

When it comes to estimating the magnitude of genetic drift, there is hardly any indexes other than the effective population size. Starting from the binomial sampling distribution, this research proposed using mean deviation of allele frequency change as a direct measurement of drift, then tested it in a classical example concerning unequal breeding sex ratio. This study found that: (1) mean deviation offers a new dimension in measuring the magnitude of drift; (2) the measurement displays a “half-half” pattern; (3) allele frequency determines the efficacy of hitchhiking effect of rare alleles, and in what way that “half-half” pattern should be divided.

## Introduction

As one of the fundamental evolutionary forces, genetic drift has been explored using models like random walk and coalescent process. The former looks forward in time, measuring drift by following the fluctuation of allele frequency—bigger step size translates into more significant drift; the latter looks backward in time^[1]^, measuring drift by effective population size (*N_e_*)–smaller *N_e_* corresponds to stronger drift^[2, 5]^. Another approach, the Wright-Fisher model, treats genetic drift as a binomial sampling process, and becomes the most common model to study drift^[4]^. Started from binomial sampling, this research incorporated both allele frequency change (i.e., step size in random walk model) and *N_e_* (key component in coalescent theory), and proposed a direct and exhaustive method to measure the magnitude of drift.

In terms of allele frequency, drift is by definition the change in allele frequency due to random sampling, and binomial distribution provides the probability of this change, so the combination of these two factors gives the mean deviation (*MD*), a direct measurement of drift (see below).

In terms of *N_e_*, breeding males and females (*N_m_* and *N_f_*, respectively) in an idealized population are the same, but unequal sex ratio reduces *N_e_* and intensifies genetic drift^[6]^ by:

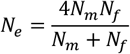

For instance, a population where 1 male breeding with many females would experience genetic drift as strong as an idealized population of 4. And a 2-male population would withstand drift as an idealized population of 8 does, and 3-male as another one of 12 does, etc. Classical coalescent theory treats alleles as identity by descent, so their frequencies are left out from this equation. This research introduced allele frequency into the calculation of the magnitude of drift, for populations of 1, 2, and 3 males breeding with many females, then compared the results with the idealized populations.

## Methods

### 1. Magnitude of drift

In a neutral model of one locus and two alleles (A_1_, A_2_), binomial distribution gives the probability of randomly sampling *k* A_1_ into the next generation (containing 2*N* alleles), based on A_1_ frequency (*p*) in the current generation:

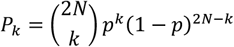

Considering the absolute change in allele frequency is |*k*/2*N*–*p*|, then by definition, the average magnitude of drift can be written as the weighted mean deviation (*MD*) of allele frequency change:

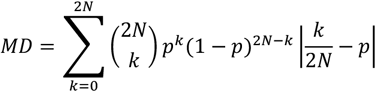

### 2. *MD* in extremely female-biased population where only 1 male breeds

In this case, *N_e_*≈4, so there are 7 values of interest for *p* (*p*=0/8 and *p*=8/8 are not meaningful to discuss with respect to drift).

Under no selection and constant population size, when forming the next generation, *N* gametes of each sex would be sampled. Each of the two alleles from that single male is expected to drift to *N*/2 copies, while each allele from females has only about 1/2 chance to reach the next generation.

For that male individual, the gametes are generated through a process of sampling with replacement. Denote *x* as the number of A_1_ allele(s) in each of the 3 possible genotypes (hereafter categories of A_1_ number in breeding males, see below), and *m* as the numbers of A_1_ alleles entering the next generation from that male, then the probability of sampling *m* A_1_ alleles from that male (*P_m_*) can be written as:

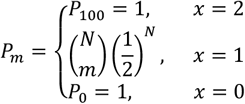

The number of A_1_ in breeding females is subject to that in breeding males because the total has to sum up to 2*Np*, so the sampling process (*N* from 2*N*–2 alleles) is without replacement. Thus, the probability of sampling *f* A_1_ alleles from female (*P_f_*) follows hypergeometric distribution:

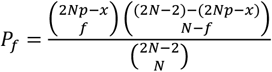

The change in *p* is:

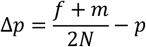

Then generally, the *MD* of each genotype can be calculated through:

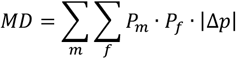

The following example concerns a population of 99 females and 1 male, and the population size remains constant (*N*=100).

#### 2.1 *x*=2 (A_1_A_1_)

In this category, all the 100 sperms contain A_1_, so *m*=100 and *P_m_*= *P*_100_=1, thus,

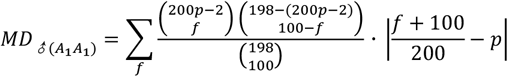

#### 2.2 *x*=1 (A_1_A_2_)

In this category, the male gamete pool is sampled 100 times and the probability of getting A_1_ is ½ for each sample, thus,

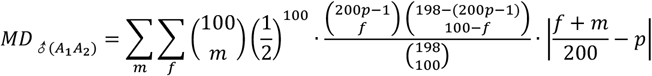

#### 2.3 *x*=0 (A_2_A_2_)

In this category, none of the 100 sperms contains A_1_, so *m*=0 and *P_m_*= *P*_0_=1, thus,

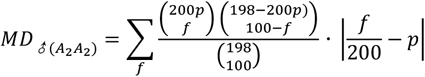

#### 2.4 Weighted *MD*

Because the parental population are of fixed size and allele frequencies, determining which two gametes get into that male is also a sampling process without replacement. Thus, the genotype frequency can be calculated through 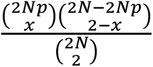, and the *MD* weighted by genotype frequencies is:

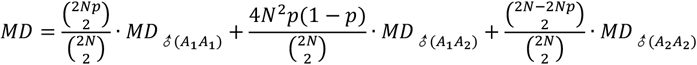

### 3. More than 1 male breeds

As long as the number of females is far more than males, it does not affect the essence of the result (data not shown). To avoid interruption during the binomial sampling due to non-integers, this research set the population size to 1000 for the 2-male case, and to 600 for the 3-male case.

#### 3.1 2 males breed with 998 females

In this case *N_e_*≈8, meaning that there are 15 meaningful values for *p*. In these 2 males there are 4 slots for alleles to fill, so *x* (number of A_1_ in both males) ranges from 0 to 4, corresponding to 5 categories, then we have:

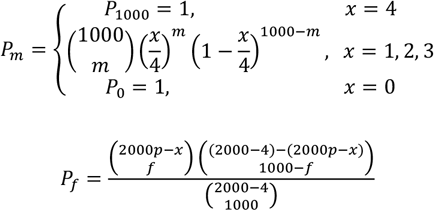

The weight for each category is:

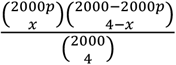

Other calculations are in the same procedure as the 1-male case.

#### 3.2 3 males breed with 597 females

In this case *N_e_*≈12, meaning that there are 23 meaningful values for *p*. In these 3 males there are 6 slots for alleles to fill, so *x* (number of A_1_ in all 3 males) ranges from 0 to 6, corresponding to 7 categories, then we have:

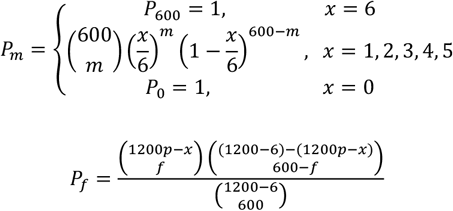

The weight for each category is:

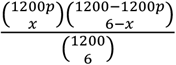

Other calculations are in the same procedure as the 1-male case.

All *MDs* were calculated with R^[3]^.

## Results and discussion

### 1. 1-male case

Table 1 shows that, with *MD* as an index of drift, the two populations of the same *N_e_* can still experience slightly different degrees of genetic drift, and when *p*=1/2 the two populations are the closest.

**Table 1.** *MDs* of the 1-male case.

| *MD* | *A*<sub>1</sub> frequency (*p*) |  |  |  |  |  |  |
| --- | --- | --- | --- | --- | --- | --- | --- |
|  | 1/8 | 2/8 | 3/8 | 4/8 | 5/8 | 6/8 | 7/8 |
| idealized (*N<sub>e</sub>*=4) | 0.08590223 | 0.1167984 | 0.132015 | 0.1367188 | 0.132015 | 0.1167984 | 0.08590223 |
| ♀biased (*N<sub>e</sub>*≈4) | 0.09466843 | 0.1389715 | 0.1447774 | 0.1353594 | 0.1447774 | 0.13897142 | 0.094668416 |
| ♂(*A*<sub>1</sub>*A*<sub>1</sub>) | 0.4330808 | 0.3712121 | 0.3093434 | 0.2474747 | 0.1856061 | 0.1237374 | 0.06186869 |
|  | **0.006528856** | **0.022851** | **0.04313707** | **0.06155778** | **0.07228377** | **0.06948569** | **0.04733421** |
| ♂(*A*<sub>1</sub>*A*<sub>2</sub>) | 0.1856061 | 0.1237375 | 0.06231416 | 0.02436533 | 0.06231416 | 0.1237375 | 0.1856061 |
|  | **0.04080536** | **0.0466347** | **0.02935654** | **0.01224388** | **0.02935654** | **0.04663473** | **0.04080535** |
| ♂(*A*<sub>2</sub>*A*<sub>2</sub>) | 0.06186869 | 0.1237374 | 0.1856061 | 0.2474747 | 0.3093434 | 0.3712121 | 0.4330808 |
|  | **0.04733421** | **0.06948576** | **0.07228378** | **0.06155778** | **0.04313709** | **0.022851** | **0.006528856** |

The *MDs* of female-biased population are subdivided according to the male's genotypes, and weighted values are in bold.

Besides, subdividing the *MD* of the female-biased population allows us to zoom in for more detail. Traditionally, this classical case has been used to illustrate how a male’s rare allele can drift a big step by luck, but the low probability for having this allele could largely cancel out this effect. Take *p*=1/8 for example, when the male is A_1_A_1_, every offspring would carry A_1_ and *p* would have a dramatic increase, translating to a very high *MD* (>0.4). However, since A_1_ is a fairly rare allele (*p*=1/8), the chance to have an A_1_A_1_ male is even lower, so the weighted *MD* is about 0.0065, which does not contribute much to the total *MD* (Fig. 1A).

**Fig. 1.**
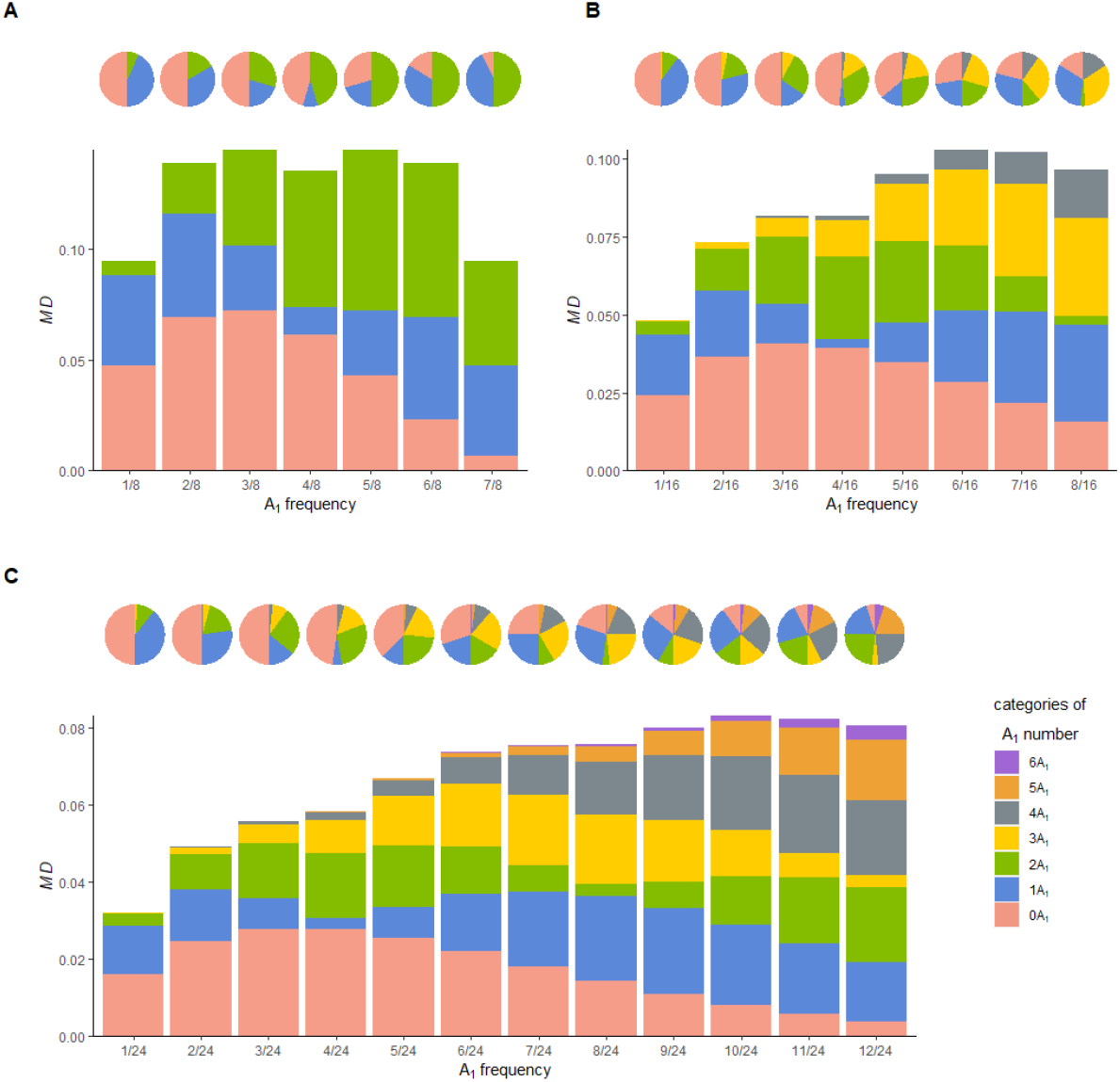
Weighted *MDs* at various frequencies of A_1_ for 1-male (A), 2-male (B), and 3-male (C) cases. For B and C, the second half is omitted due to symmetry. *MDs* are subdivided according to categories of A_1_ number in the breeding male(s), and these categories always form a “half-half” pattern (the pie charts).

A common concern about highly biased breeding sex ratio is that deleterious alleles could soar in number, but this is the case only if those alleles can find their place in the breeding male, which is not very likely. Even if the male is selected to breed for traits on certain loci, hitchhiking alleles on other independent neutral loci are sampled randomly based on their frequencies. In other words, biased sex ratio alone is not enough to efficiently boost up a rare allele in number.

### 2. 2-male and 3-male cases

Consistent with the 1-male case, two types of population have similar but not the same *MDs,* which are closest to each other at *p*=1/2 (Table 2, 3).

**Table 2.**
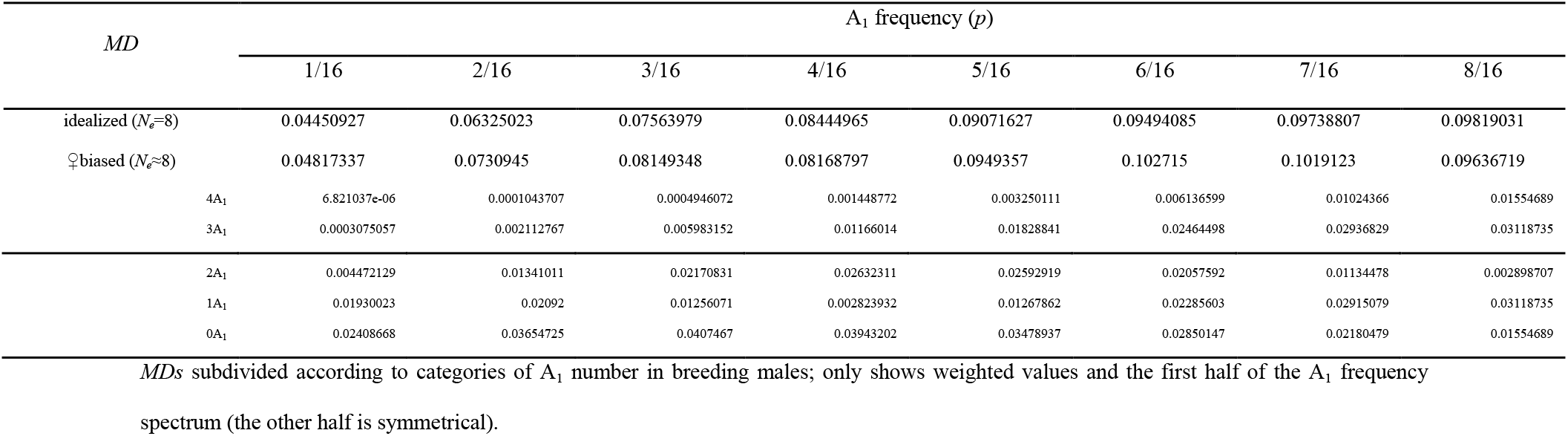
*MDs* of the 2-male case.

**Table 3.** *MDs* of the 3-male case.

| *MD* | $A_1$ frequency ($p$) | | | | | | | | | | | |
| --- | --- | --- | --- | --- | --- | --- | --- | --- | --- | --- | --- | --- |
|  | 1/24 | 2/24 | 3/24 | 4/24 | 5/24 | 6/24 | 7/24 | 8/24 | 9/24 | 10/24 | 11/24 | 12/24 |
| idealized ($N_e=12$) | 0.03000662 | 0.04317759 | 0.052367 | 0.05940697 | 0.06499195 | 0.06947143 | 0.07304815 | 0.07584952 | 0.07795895 | 0.07943146 | 0.08030203 | 0.08059013 |
| ♀biased ($N_e \approx 12$) | 0.032096942 | 0.04913652 | 0.055738827 | 0.058387799 | 0.067052887 | 0.073641601 | 0.075413233 | 0.075709863 | 0.080016574 | 0.083166919 | 0.082440734 | 0.080659188 |
| 6$A_1$ | 1.849796e-09 | 1.327306e-07 | 1.519557e-06 | 8.341719e-06 | 3.069468e-05 | 8.771458e-05 | 0.0002104102 | 0.0004436486 | 0.0008467416 | 0.001492078 | 0.00246125 | 0.003838119 |
| 5$A_1$ | 2.343074e-07 | 7.544684e-06 | 5.34465e-05 | 0.0002053346 | 0.0005637799 | 0.001248818 | 0.00237855 | 0.00404338 | 0.006279208 | 0.009042898 | 0.01219335 | 0.01548149 |
| 4$A_1$ | 1.155117e-05 | 0.0001679435 | 0.0007341357 | 0.001962334 | 0.00398733 | 0.006772968 | 0.01010143 | 0.01359706 | 0.01677634 | 0.01911588 | 0.02012995 | 0.01944927 |
| 3$A_1$ | 0.0002758736 | 0.001810524 | 0.004831236 | 0.008836589 | 0.01298487 | 0.01638283 | 0.01828587 | 0.01822074 | 0.0160418 | 0.01193272 | 0.006512489 | 0.00312143 |
| 2$A_1$ | 0.003146291 | 0.009120055 | 0.01424072 | 0.01668576 | 0.01593751 | 0.01232856 | 0.006784793 | 0.003100204 | 0.006815464 | 0.01252548 | 0.01689294 | 0.01944927 |
| 1$A_1$ | 0.01261452 | 0.01346206 | 0.008018969 | 0.00299108 | 0.008117812 | 0.01477756 | 0.01941946 | 0.02183749 | 0.02222073 | 0.02096359 | 0.01855249 | 0.01548149 |
| 0$A_1$ | 0.01604847 | 0.02456826 | 0.0278588 | 0.02769836 | 0.02543089 | 0.02204315 | 0.01823272 | 0.01446734 | 0.01103629 | 0.008094273 | 0.005698265 | 0.003838119 |

*MDs* subdivided according to categories of $A_1$ number in breeding males; only shows weighted values and the first half of the $A_1$ frequency spectrum (the other half is symmetrical).

Combining the 1-male case, an interesting pattern emerges: the total *MD* splits in half, each of which is composed of certain categories of A_1_ in breeding males, and the composition varies with A_1_ frequency. Take the 3-male case as an example: from *p*=1/24-3/24, half of the total *MD* comes from 0A_1_ category, and when *p*=4/24, the weighted *MD* of 0A_1_ category plus 1/2 of that from 1A_1_ category makes up exactly ½ of the total *MD*. This pattern carries on for all the subsequent frequencies: from *p=5/24-7/24,* the sum of 0A_1_ and 1A_1_ equals ½ of the total *MD,* and when *p* reaches 8/24, an extra ½ of 2A_1_’s *MD* is needed to maintain this “half-half” pattern, and same cycle is repeated afterwards (Table 3, Fig. 1C).

The exact pattern is detected in 1-male and 2-male cases (Table 1-2, Fig. 1A-B), and it carries on beyond (e.g., 4-male, data not shown). A chart (Fig. 2) illustrates this pattern for one to refer to: it is the abstract representation of a table, and one can locate the boundary of “half of the drift” for a certain A_1_ frequency and number of breeding males.

**Fig. 2.**
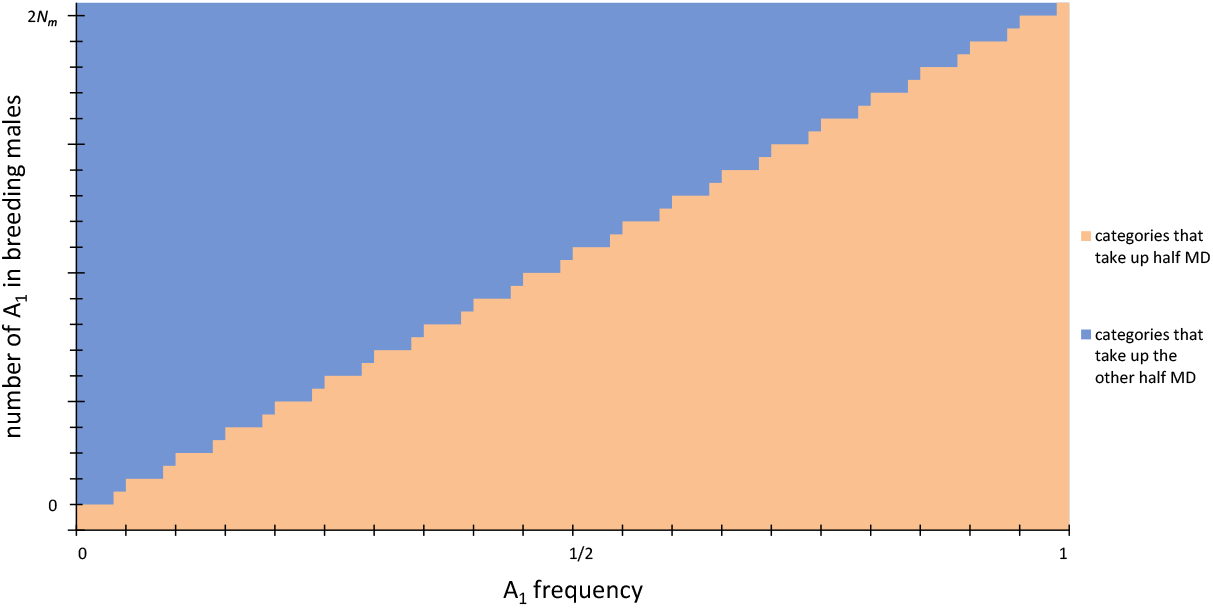
Generalization of the “half-half” pattern. The nesting zigzag shape is to show that half or all of the *MD* from the following allele frequency level are added periodically.

Previous studies on coalescent theory and *N_e_* have discussed that, random chance experienced by a diploid organism comes from two sources: the first is within individual, acting when an allele gets into the successful gamete during Mendelian Segregation; and the second is between individuals, acting during survival and reproduction. Genetic drift can result from both, and under some peculiar circumstances, these two sources of randomness turn out to be equal^[4]^.

This research found that the magnitude of drift is divisible in another aspect (category of A_1_ number) for populations of extremely biased sex ratio. The division helps in understanding the property of drift: as illustrated in Fig. 2, how to break up the *MD* is a function of the number of breeding males and allele frequency on that locus; when variation of allele frequency is very small (*p* close to 0 or 1), those two halves of the *MD* are distributed extremely unevenly. This finding could be of especial interest when studying the impact of neutral drift on rare alleles.

## Conclusions

When measuring drift in a population with unequal breeding sex ratio, an idealized population and a female-biased population consistently have similar *MDs*; for the latter, *MD* can always be divided into two equal parts, each of which regularly contains certain categories of A_1_ number in breeding males; a rare neutral allele can hardly multiply through hitchhiking, because it requires a great deal of luck to end up in the breeding males. In the future, it would be interesting to test this method in another classical case: fluctuating population size over generations. Besides, incorporating selection, migration and directionality in frequency change are also worth of pursuing.

## Acknowledgement

I thank Sean Rice for insightful discussion, Meijun Dong for helps in some graphs, and Maosheng Zhang for suggestion in simplifying some equations.

## References

[1] Kingman, J.F.C. (2001). Origins of the Coalescent: 1974-1982. Genetics, 156(4), 1461–1463.

[2] Nomura, T. (2003). Effective size of populations with heritable variation in fitness. Heredity. 89, 413–416. 10.1038/sj.hdy.6800169.

[3] R Core Team. (2021). R: A language and environment for statistical computing. R Foundation for Statistical Computing, Vienna, Austria. URL https://www.R-project.org/.

[4] Rice, S. (2004). Evolutionary theory: mathematical and conceptual foundations. Sunderland, MA: Sinauer Associates, 73–117.

[5] Wang, J., Santiago, E., & Caballero, A. (2016). Prediction and estimation of effective population size. Heredity, 117(4), 193–206. https://doi.org/10.1038/hdy.2016.43

[6] Wright, S. (1931). Evolution in Mendelian populations. Genetics, 16, 97–159.

